# ScreenGarden: A shinyR application for fast and easy analysis of plate-based high-throughput screens

**DOI:** 10.1101/2021.05.10.443457

**Authors:** Cinzia Klemm, Rowan M. S. Howell, Peter H. Thorpe

## Abstract

Colony growth on solid media is a simple and effective measure for high-throughput genomic experiments such as yeast-two hybrid, Synthetic Genetic Arrays and Synthetic Physical Interaction screens. The development of robotic pinning tools has facilitated the experimental design of these assays, and different imaging software can be used to automatically measure colony sizes on plates. However, comparison to control plates and statistical data analysis is often laborious and pinning issues or plate specific growth effects can lead to the detection of false positive growth defects. We have developed ScreenGarden, a shinyR application, to enable easy, quick and robust data analysis of plate-based high throughput assays.

## Introduction

The budding yeast *Saccharomyces cerevisiae* is a widely used model organism to understand basic molecular processes in eukaryotic cells. Over the past decades, the development of new genetic techniques enabled the creation of comprehensive clone and gene deletion libraries in yeast. These libraries can be used for many different high-throughput experiments, such as synthetic lethality and synthetic dose lethality screens^1–3^, chemical genetic screens^3^, and yeast two hybrid and Synthetic Physical Interaction screens to unravel unknown protein-protein interactions^4,5^. Although investigating different research aspects, the common read-out of these screening methods is colony growth on solid media. Libraries are typically organised in arrays of 96, 384 and 1536 colonies per plate and the colony size of experimental and control conditions are compared to determine growth effects. Library based screens efficiently generate robust and large datasets in a short time, however, data analysis can be challenging and merely visual comparison of colonies lacks normalisation of plate differences and is highly subjective. Quantitative data analysis of colony growth can be used to define the strength of growth defects in an unbiased manner. Colony size can be quantified using tools such as ImageJ, HT Colony Grid Analyser, CellProfiler, gittr or Spotsizer^6–9^, but these have limited capacity for downstream analysis. Other tools which allow more sophisticated data analysis include proprietary tools such as PhenoBooth Colony Imager (Singer Instruments Ltd, UK), or the *ScreenMill* software suite. The ‘DR engine’ and ‘SV engine’ of the *ScreenMill* software suite were developed to facilitate statistical analysis and offered web-based applications which allowed reviewing and visualising of screening data^6^. However, *ScreenMill* was designed to compare each experimental plate to a single control. Assays that compare experimental plates to two controls, such as Synthetic Physical Interaction screens, require further data processing, which is laborious and can lead to errors. Furthermore, the *ScreenMill* web application is currently not accessible and the software is composed of different programming languages making it difficult to run the analysis for inexperienced data analysts.

To simplify the analysis of plate-based high throughput screens, we have developed ScreenGarden, a Shiny R application (https://shiny.rstudio.com/) for statistical analysis of screen-based assays independent of *ScreenMills*’ ‘DR engine’ and ‘SV engine’. ScreenGarden can be run as a web application or offline using RStudio (https://www.rstudio.com/), which makes it also very easy to adjust and customise the script. ScreenGarden is further developed to facilitate screen analysis which compare experimental plates to two independent controls. Furthermore, ScreenGarden allows direct quality control of screens and plotting of data without exporting the output files into other data analysis software. At the same time, ScreenGarden produces the raw numbers behind each step of data analysis enabling more sophisticated data interpretation downstream for users who require this.

## Results

The ScreenGarden application was designed to enable statistical analysis of plate-based assays using colony size as a readout. Data analysis using ScreenGarden can be performed in one step if there is a single control condition for each experiment (Figure 1). Log growth ratios (LGRs) and Z-scores are calculated for a single control using the ‘CalculateLGRs’ command. The user can download a ‘mean file’ which averages the data over the number of replicates, or a ‘replicates file’, which contains the separate data of each individual replicate. Additionally, ScreenGarden can combine data from experiments that use two independent controls using the ‘Combine2controls’; the comparisons to two different controls are combined and downloaded as a ‘merge file’. Finally, the data can be plotted automatically for analysis and quality control, and plots can be directly downloaded from the website. A detailed description of how to use the ScreenGarden web application can be found in the appendix (Supplementary file 1) or downloaded from the ScreenGarden homepage. https://screengarden.shinyapps.io/screengardenapp/

**Figure 1.**
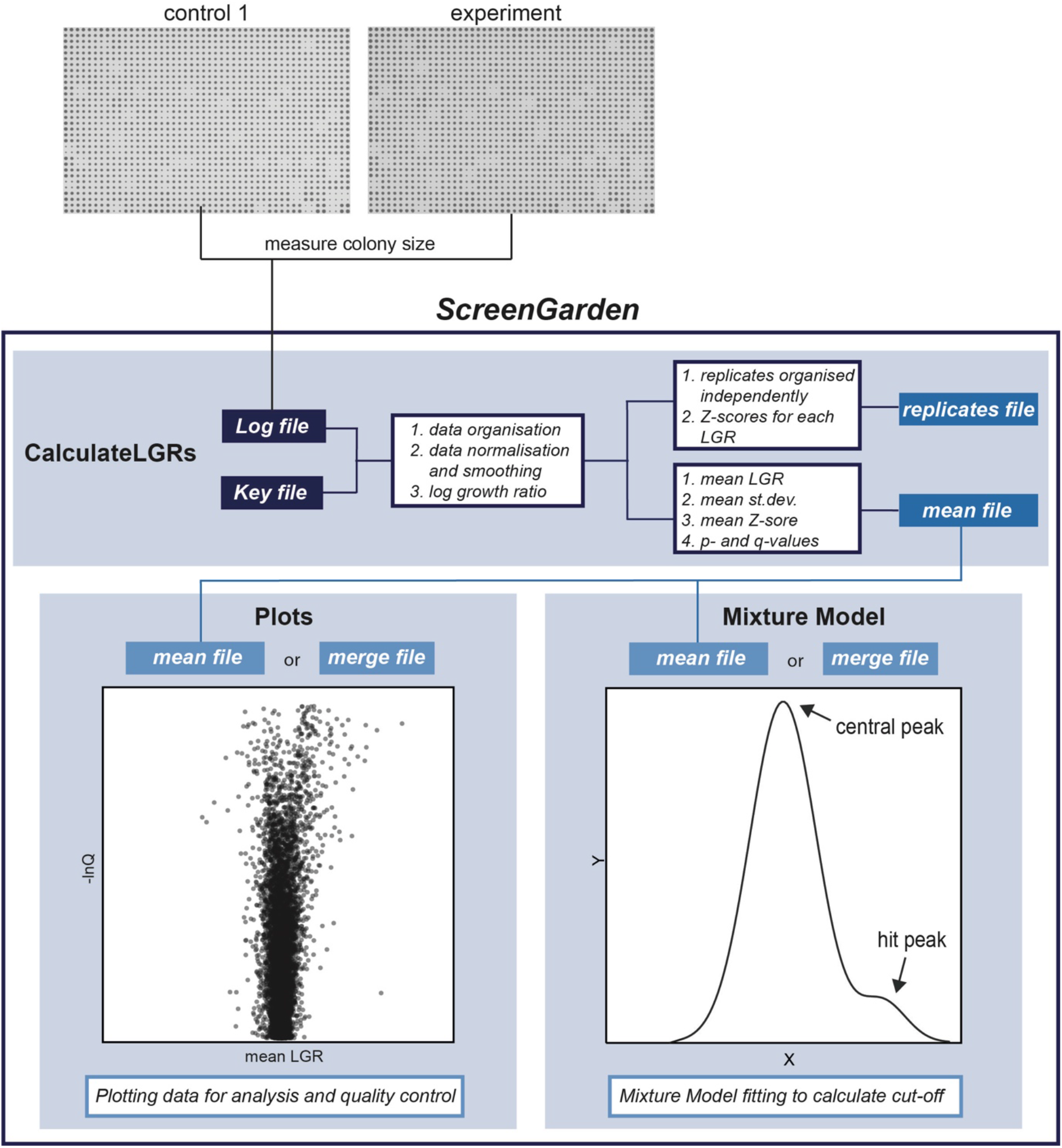
Steps of data analysis using ScreenGarden. The ScreenGarden application offers a tool for stepwise analysis of plate-based high-throughput screens and was specifically adapted to facilitate the analysis of screens with one or two independent controls. The ‘Calculate-LGRs’ script performs statistical analysis of colony sizes compared to a single control. Using the ‘Combine2Controls’ script, two comparisons can be combined to generate a ‘merge file’ with average LGRs and Z-scores. At last, the data can be uploaded and plotted for quality control, data analysis and data visualisation.

Here, we applied ScreenGarden analysis to previously published data from genome wide Synthetic Physical Interaction screens^10^ to compare this software with another established method. Synthetic Physical Interaction screens rely on a GBP-GFP binding system to forcibly associate GPB-tagged proteins to the yeast proteome^5^ and a negative impact on cell-growth upon protein-protein interaction is defined as a Synthetic Physical Interaction.

### Comparison of experimental and control colony sizes

ScreenGarden analysis can be performed with different array sizes (384 and 1536 colonies on each plate) and a replicate number of 4 or 16 colonies per yeast library strain. The software requires a ‘Log file’ and a ‘Key file’ as input files. Here, we used *ScreenMills’ CM Engine* to measure colony sizes on plate, which automatically produces a ‘Log file’ as a list of colony sizes ordered by plate position (starting from A1, A2, A3 … H12 for 96 colonies on plate, Supplementary file 2). Other software tools, such as HT Colony Grid Analyser, can be used to measure colony sizes on plate, but files have to be converted to the specified format (Supplementary file 3). The ‘Key file’ contains information about the yeast library and assigns the genotype of each strain to its specific plate position (Supplementary file 4). It is necessary that both, ‘Log file’ and ‘Key file’ are in the format as shown in the examples and that the files are uploaded into ScreenGarden as tab delimited or comma separated files. Here, we applied ScreenGarden analysis to Synthetic Physical Interaction screen data with the outer kinetochore subunit Dad2 (Supplementary data 1). In this screen, GBP-tagged Dad2 was recruited to 6234 different GFP-strains and the screen was performed in 4 replicates, 1536 colonies on a total of 17 plates. First, colony sizes are normalised by the plate median to correct for plate specific effects on growth. Median plate correction is important to prevent false-positive growth defects which might occur due to differences in nutrition, humidity and other external factors^11^ (Figure 2A,B). However, for screens with a high number of growth defects, median-normalisation should be omitted, as a low median colony size might reflect an experimentally valid negative effect on growth. Alternatively, the data can be normalised to the median growth value of positive control colonies located at specific positions on the plate which have to be identified as ‘Control’ in the open reading frame (ORF) defining column in the ‘Key file’. A second difficulty for plate-based screens are spatial anomalies within a plate array. Colonies often grow faster at the plate periphery (Figure 2C) as there is less competition for nutrients^12,13^. We have incorporated a simple smoothing algorithm from Ólafsson and Thorpe^5^, which adjusts colony sizes based on their plate position to counteract spatial anomalies across the plate. Incorporating the smoothing algorithm into ScreenGarden analysis successfully limits spatial effects (Figure 2F). After median-correction and smoothing, LGRs are calculated separately for each replicate of each strain on each plate. The LGR is the natural logarithm of the ratio of the control colony size divided by the experimental colony size. The difference between the control and experimental replicates is evaluated by applying a Student’s t-Test to generate a p-value. To compensate for false positive growth defects, which naturally occur in large-scale screens, these p-values are adjusted using a false discovery rate (FDR) correction method after Benjamini and Hochberg ^14^, resulting in more conservative q-values. For both, p-values and q-values, the negative natural logarithm is determined, which is useful for generating Volcano plots which compare LGRs against their p- or q-values. Subsequently, mean LGRs are determined as the average of the 4 or 16 replicate LGRs. Finally, Z-scores for each mean LGR are calculated, which can be used to assess growth defects.

**Figure 2:**
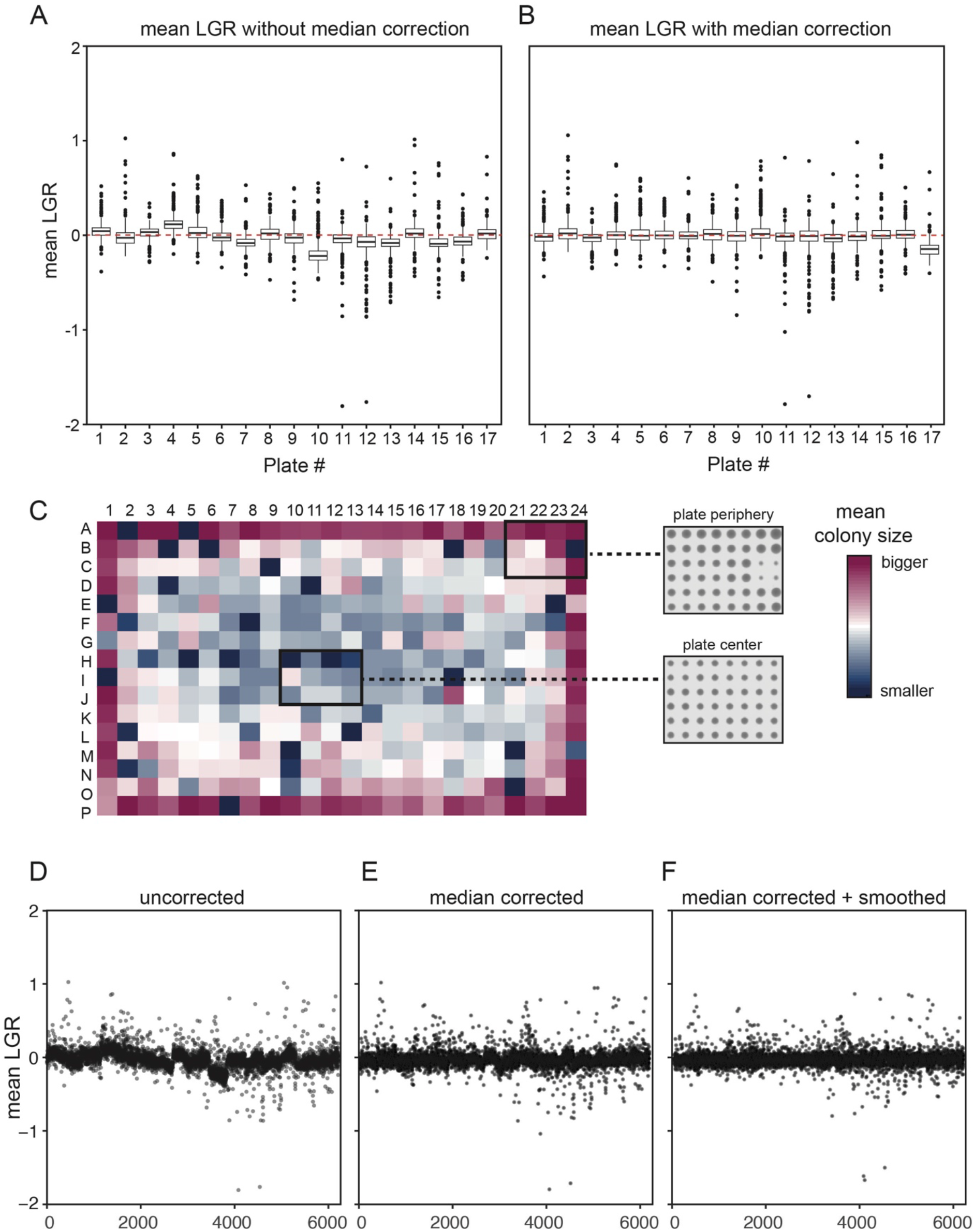
Median-correction of colony sizes reduces the influence of plate differences. (A)External factors and discrepancies in pinning can result in differences in colony growth between plates. The mean LGRs organised by plate without median correction are plotted. (B) Correction of colony sizes using the plate median counteracts these plate differences, median-correction data from (A) is plotted. (C) A heatmap shows spatial anomalies of colony sizes on plate especially at the plate periphery. Red squares indicate a colony size greater than the plate median and blue squares highlight smaller colonies. The inset shows an example of the raw data. (D-F) Mean LGRs are plotted against the yeast library with the data organised on the x-axis, by plate, row and column to highlight the impact of plate differences and spatial anomalies. Including the spatial smoothing algorithm further abolishes any differences based on colony position on plate.

The ‘CalculateLGRs’ tool produces two output files, a ‘replicates file’ where each replicate is listed independently, and a ’mean file’ which contains the averaged LGRs and Z-scores of the replicates combined. Both files can be downloaded directly from the website as ‘.csv’ files and easily imported into R, Excel and other applications for data analysis.

### Combining two independent controls

The second, optional step of screen analysis using ScreenGarden is the ‘Combine2controls’ tool, which is designed for plate-based screens with two independent controls (Figure 3A). After separately running the ‘Calculate LGRs’ script with each control, the resulting two ‘mean files’ can be uploaded and joined to a single ‘merge file’. The ‘merge file’ includes all the information from the single control analyses and further includes the mean LGRs and Z-scores and the maximum of p- and q-values from both controls. We chose the maximum p or q-value from both independent control datasets as measure for significance rather than calculating combined p- and q-values using Fisher’s method^15^, since the data originates from a single experimental dataset with different controls and the two p and q values are not truly independent. The maximum q-value should not be considered a measure of statistical likelihood for mean LGRs of two controls, but rather facilitate the identification of false positive growth defects based on pinning errors. The ‘Plot’ function of ScreenGarden allows these data to be compared, for example to compare the LGR values produced by the two controls. For example, for a Dad2 Synthetic Physical Interaction screen dataset, 18.2% (control 1) or 29.2% (control 2) of observed growth defects from single control comparisons were excluded using the average LGR (Figure 3B,C). Hence, ScreenGarden automatically defines a set of high confidence growth defects for screens based upon two independent controls.

**Figure 3:**
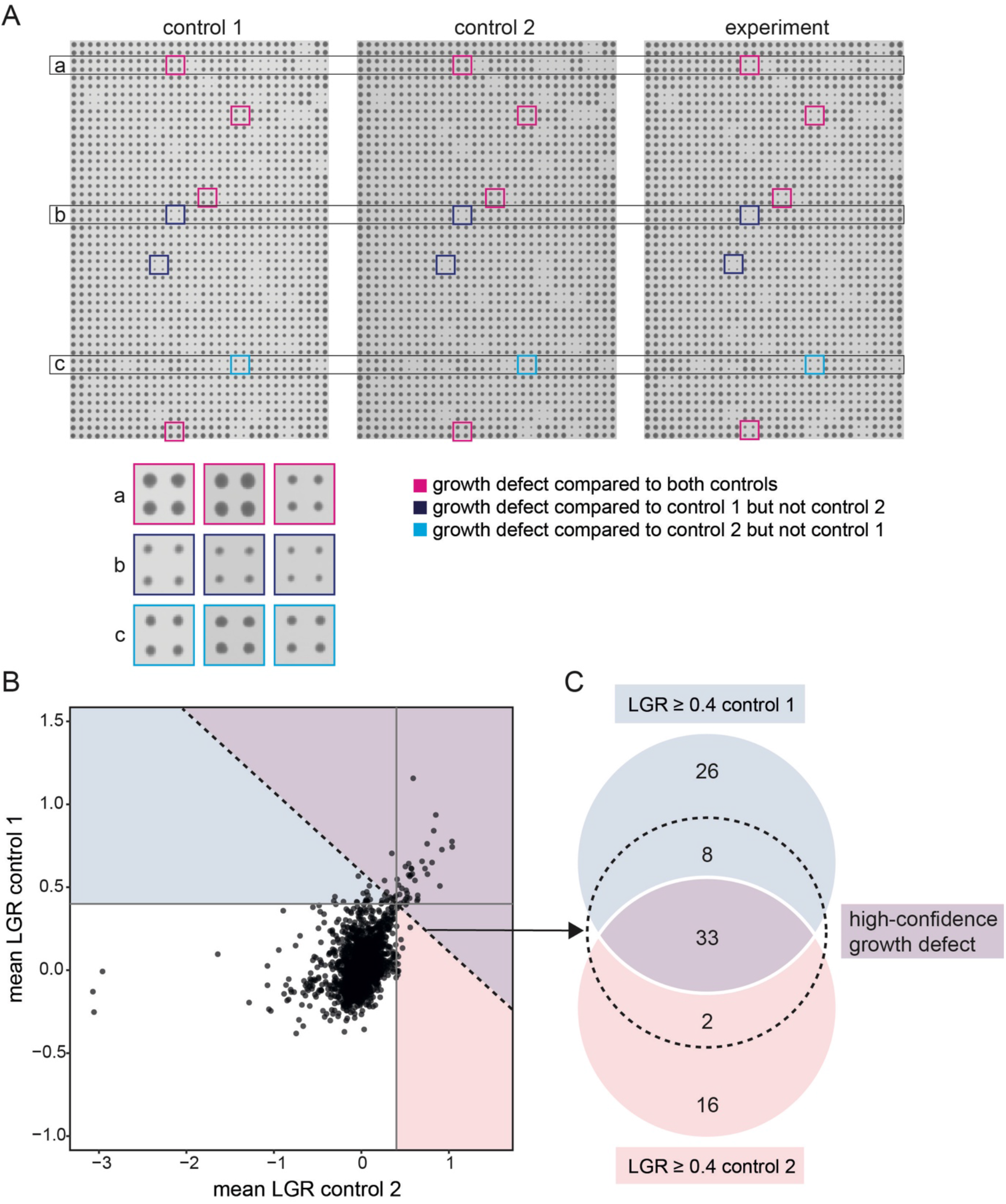
Combining analyses with two independent controls using ScreenGarden. (A) The raw data (plate images) from two control plates to the same experimental plate are shown together for one 1536 density plate with four replicates per strain. Control-specific hits are highlighted in red (control 1) and blue (control 2) boxes respectively. (B) The LGR values for comparing the experiment independently with each control are plotted. The dashed line visualises the empirical cutoff value of LGR ≥ 0.4 for the average LGRs of both control comparisons. (C) These data are shown as a Venn diagram, with the dashed circle indicating all data with an average LGR (of the two controls) ≥ 0.4. Using two controls rather than one defines a subset of high-confidence growth defects and excludes control-specific effects.

### Quality control using ScreenGarden

ScreenGarden can be used to plot results directly without laborious reimporting and reformatting in a different application, such as R or Excel. Using the ‘Plots’ tab, either the ‘mean file’, if experiments are compared to one control, or the ‘merge file’, if the screen was performed using two controls, can be uploaded and any two columns can be plotted against each other. Plotting is useful for quality control of screen data, and for example, it allows users to assess the data plate by plate to identify whether any plates produced anomalous LGR values (Figure 4A). Since the data can be plotted by Row or Column, the data can be scrutinised to ask whether the smoothing algorithm efficiently reduced spatial effects, i.e. whether or not specific rows or columns have higher or lower LGR values (Figure 4B,C). In the ‘Plot’ function the distribution of LGRs is automatically visualised in a histogram with an adjustable number of bins (Figure 4D). Plotting mean LGRs against the negative natural logarithm of p- or q-vales respectively allows for rapid assessment of reproducibility, as high p-/q-values account for a large difference in replicate colony sizes. Hence, strains that have inconsistent replicates in the data can be easily identified and if necessary excluded from further analysis. Only three Synthetic Physical Interactions of the Dad2 dataset were above the q-value threshold of 0.05. We used ScreenGarden to analyse Synthetic Physical Interactions with the nucleolar protein Nop10, a second dataset from Berry and colleagues^10^ (Supplementary data 2). Nop10 association caused a higher number of growth defects compared to the Dad2 dataset, and we found 19 growth defects with a q-value below the threshold (Figure 4F). Notably, strains with a low value for -lnQ vary in replicate colony sizes (Figure 4G). We compared these growth defects to the validation screens performed by Berry and colleagues, who identified 7 of these 19 interactions as false positive growth defects (Figure 4G,H). Exclusion of growth defects based on high q-values decreased the false-discovery rate for the Nop10 screen by approximately 5% (Figure 4H).

**Figure 4:**
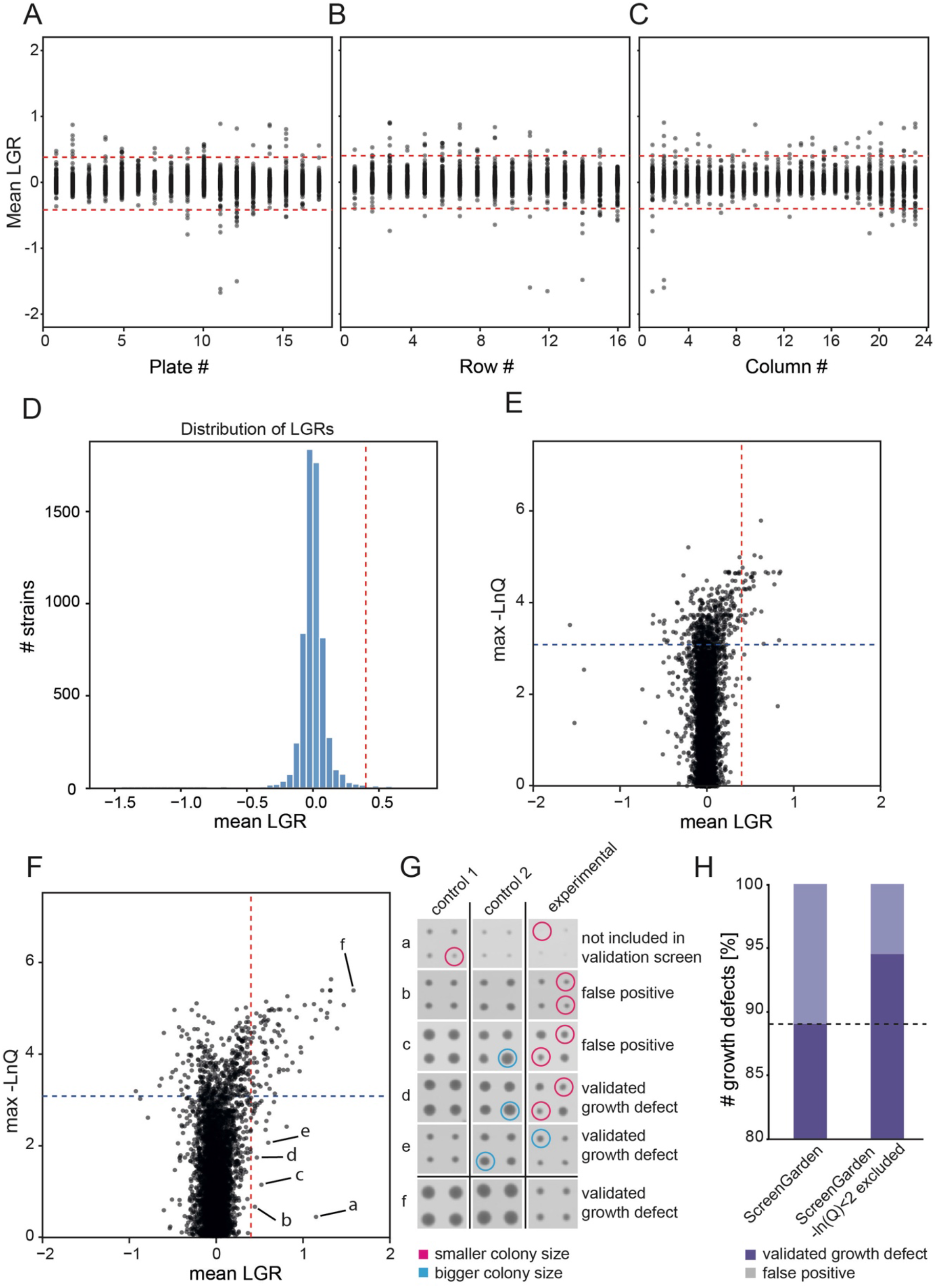
Quality control of plate-based screens using ScreenGarden. Using the ‘Plots’ tab, users can upload .csv files downloaded from ‘ClaculateLGRs’ or ‘Combine2Controls’ and plot any column against each other. For quality control, mean LGRs were plotted against plate (A), row (B) and column (C) number. (D) Histogram showing the distribution of data. The majority of LGRs are distributed close to zero. The red dashed line highlights a LGR of 0.04. (E) Negative max lnQ values were plotted against mean LGRs to identify replicate inconsitencies in the Dad2 Synthetic Physical Interactions data set. The red dashed line highlights a LGR of 0.04, the black dahed line indicates a max lnQ of 2.99 (q = 0.05). (F) Negative max lnQ values were plotted against mean LGRs to identify replicate inconsitencies in the Nop10 SPI data set. The data points labelled a to f, most with low max -lnQ values, are analysed in the next panel. (G) Selected growth defects (a to e from panel F) with a low max -lnQ value show inconsitencies in colony sizes on plate and 2 of them were identified as false positive growth defects according to Berry and colleagues^10^ (H) Exclusion based on low max -lnQ values reduced the number of false positive growth defects.

### Comparing ScreenGarden and *ScreenMill*

Next, we compared the output of ScreenGarden analysis to a previously developed tool for statistical data analysis, the *DR Engine* of the *ScreenMill* software suite (Dittmar et al., 2010) (Supplementary data 3). The *DR Engine* calculates plate median normalised LGRs but does not automatically apply a smoothing algorithm, thus we first compared unsmoothed LGRs from both *ScreenMill* and ScreenGarden (Figure 5A). As expected, the datasets are highly correlated (R^2^ > 0.99), but, notably, not identical. This observed variance might be due to *ScreenMill’s* automatic exclusion of control-dead colonies for one of the two controls. Control-dead colonies are not excluded in ScreenGarden, but plate normalised colony sizes are reported in the dataset. Since data exclusion is subjective, we allow the user to manually exclude data if the normalised control colony size is below a certain threshold (e.g. 30% of the plate median). A second explanation for the slight variation in data values of ScreenGarden compared to *ScreenMill* is the way LGRs are calculated. The LGRs are used as a measure of growth defect because if, as commonly assumed, the colony sizes are distributed according to a lognormal distribution then the LGRs will be distributed normally. *ScreenMill* calculates the LGR as ln(average control colony size/average experimental colony size) whereas ScreenGarden calculates the LGR for each colony compared to the equivalent position on the control plate before averaging across replicates of the same genotype. This latter approach of applying the logarithm before averaging is more accurate as an approximator of the mean LGR than applying the logarithm to the averaged values, since growth ratios are distributed according to a lognormal distribution and hence LGRs are distributed normally. This effect is generally small but can be significant when the variance between colony sizes is large. Next, we applied the smoothing algorithm to the unsmoothed *ScreenMill* output data and compared this to smoothed ScreenGarden data using four independent datasets (Figure 5B-E). The smoothed data correlated well for each screen (R^2^ = 0.92-0.95), however the variance was greater compared to unsmoothed data. This variance is likely based on the timepoint of smoothing. Using ScreenGarden, LGRs are smoothed before averaging and independently for each control, whereas *ScreenMill* data was smoothed manually after calculating mean LGRs of two controls. Last, we analysed the reproducibility of growth defects identified using ScreenGarden and *ScreenMill*. We compared mean LGRs ≥ 0.4 to the results of validation screens performed by Berry and colleagues to distinguish between reproducible growth defects and false positives (Figure 5F). Both ScreenGarden and *ScreenMill* performed well in identifying growth defects in the majority of screens, hence we conclude that ScreenGarden analysis can be used to successfully identify reproducible growth defects at least as effectively as *ScreenMill* analysis.

**Figure 5:**
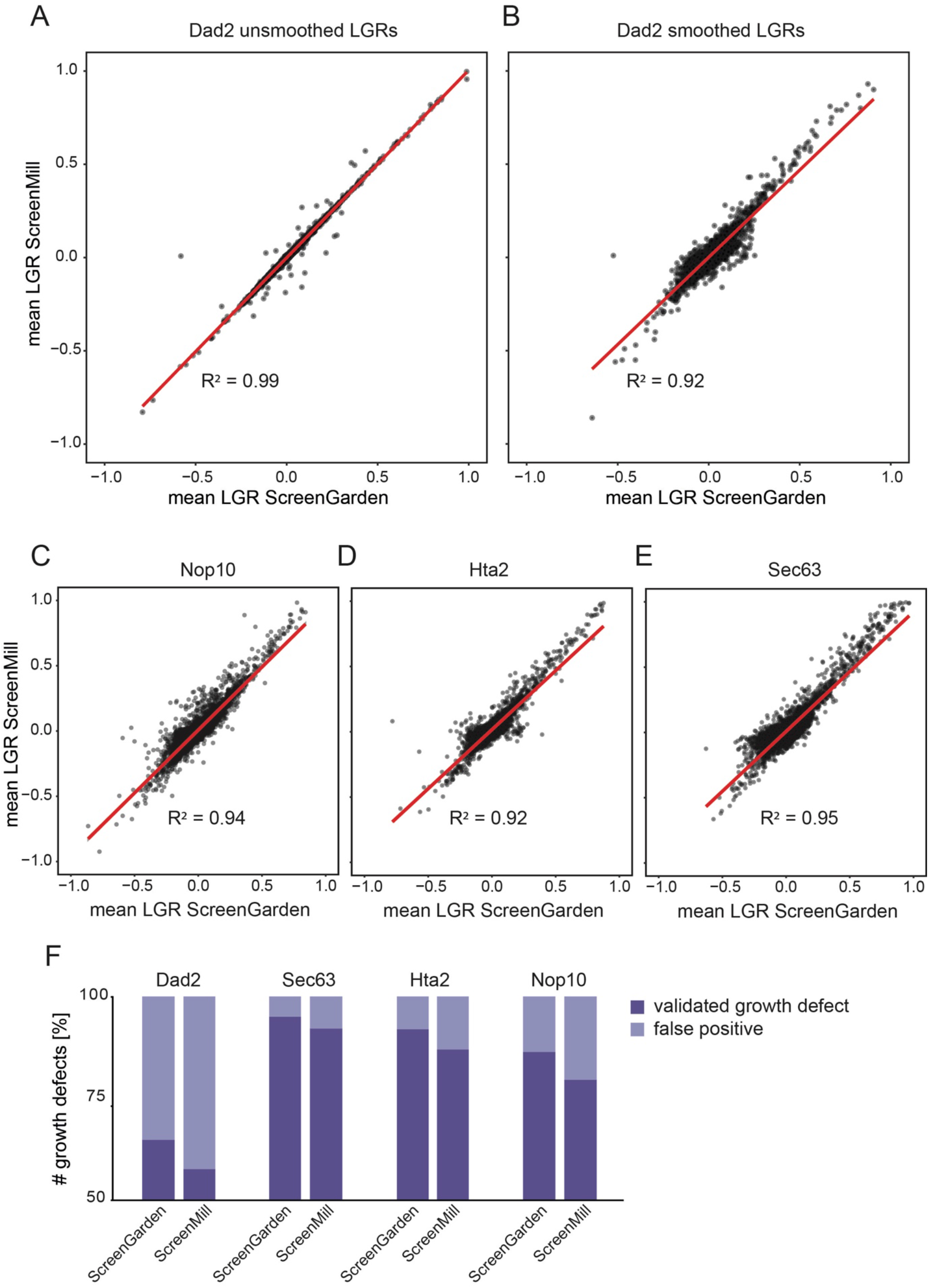
Comparison of ScreenGarden and *ScreenMill*. (A) Unsmoothed data of the Dad2 SPI screen analysed with ScreenGarden and *ScreenMill* are compared. Regression (red line) and Pearson correlation were calculated using the stats 3.6.2. package in RStudio. (B) Smoothed data of the Dad2 SPI screen analysed with ScreenGarden and *ScreenMill* are compared. *ScreenMill* does not automatically smooth data, thus smoothing was performed using PERL based on mean LGRs^5^. Smoothed data of the (C) Nop10, (D) Hta2 and (E) Sec63 SPI screens analysed with ScreenGarden and *ScreenMill* are compared. (F) Both ScreenGarden and *ScreenMill* analysis accounted for similar ratios of true growth defects and false positives when compared to the validation screen data from Berry and colleagues^10^.

### Defining cut-offs for growth inhibition

Defining the right LGR value or threshold to identify a growth effect varies from screen to screen and is often subjective. However, the threshold choice is important to prevent high rates of false positives whilst at the same time allowing sensitive identification of growth defects. In this study, we used an empirical cut-off value of LGR = 0.4 to determine growth defects, as previously defined for Synthetic Physical Interaction screens^16,17^. At LGR = 0.4, a growth defect is moderate but visible compared to control plates. However, ScreenGarden also offers mathematical approaches to define cut-off thresholds, based on the data distribution. ScreenGarden automatically calculates Z-scores, as Z-transformation fits a normal distribution to a dataset and uses the mean and variance of the data to define Z-scores for each data point. The region (−1.96,1.96) in Z-space represents the 95% of the data in a normal distribution, hence a Z-score above ^~^2 accounts for the strongest 2.5% growth effects within the data. Z-scores have an advantage of allowing datasets to be compared even when the produce quantitatively very different growth effects. However, there are several problems with using Z-scores. First, growth data are typically not normally distributed and second when a normal distribution of is applied to a large dataset, there will always be ^~^2.5% of the data with a Z-score >2 regardless of whether any growth defects were present. Screens that result in some growth defects are likely to display a multimodal or fat-tailed distribution, which are characterised by a longer tail in the positive region of the distribution curve (Figure 4D). In a previous study, Howell and colleagues have shown that proteome-wide screens such as SPI screens with a high number of growth defects can be described using bimodal normal mixture models^18–20^. Based on the mixture model, the data distribution is composed of two separate components 1 and 2 with distinct peaks (Figure 6A). Component 1 describes the central peak and contains unaffected strains with LGRs ^~^0, whereas component 2 or the ‘hit peak’ accounts for growth defects with higher LGRs. We have incorporated this script into ScreenGarden in the ‘Mixture Model’ tab, which enables the user to upload their previously calculated ‘mean file’ or ‘merge file’. The bimodal normal mixture model then calculates an FDR-adjusted q-value, with q(x) defined as the probability of inclusion in component 2, given a measured LGR of x. Hence, a q(x) = 0.5 is defined as cut-off point as LGRs are equally likely to be in component 1 or 2. We applied the mixture model fitting to SPI analysis of Nop10, as this screen resulted in a high number of growth defects (Figure 6A,B) (Supplementary data 4). We found that q ≥ 0.5 accounted for Synthetic Physical Interactions with an LGR of approximately 0.22 or higher, with a Z-score of as low as 1.3. This led to the identification of more than double the number of growth defects compared to Z-score or LGR-based cut-off definition, however, most of these additional growth defects were not included in validation screens by Berry and colleagues and thus it remains unclear if they are true growth defects or false positives (Figure 6C). Our findings suggest that using a more conservative cut-off definition, like an empirical value for LGRs when growth is visibly affected, is useful for screens without additional validation to reduce false positives. In contrast, using the bimodal mixture model and subsequent validation screening can extensively increase the number of growth defects identified in screens.

**Figure 6:**
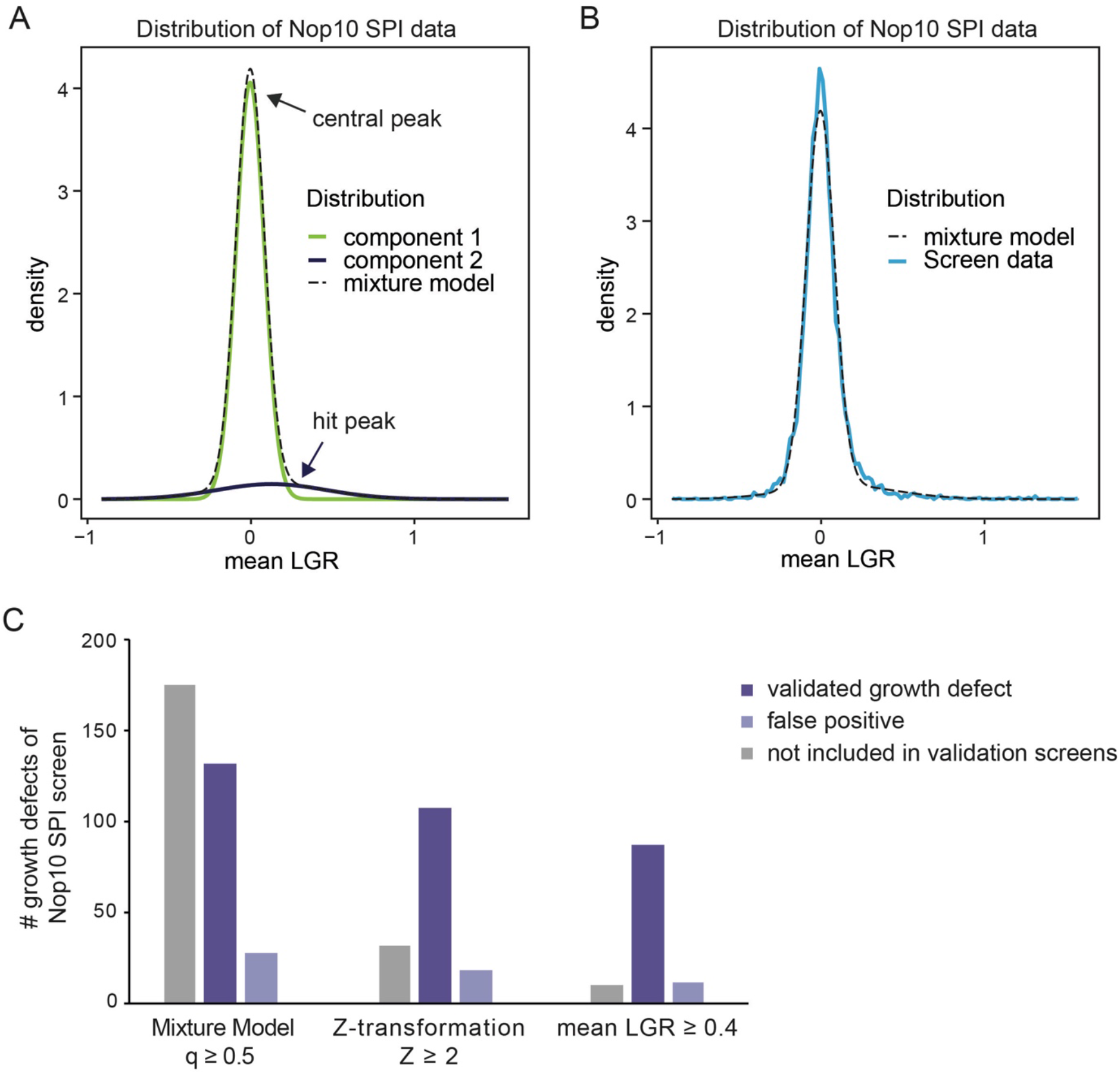
Cut-offs can be defined based on the distribution of screen data. Screens which result in a high number of growth defects are more accurately described using a bimodal mixture model and are characterised by a ‘central peak’ and a second ‘hit peak’. Bimodal mixture models can be fitted automatically to screens with many growth defects using ScreenGarden and produce a Component plot (A) and a fit plot (B) as well as q-values for each LGR. If q ≥ 0.5, the data is predicted to follow the distribution of component 2 and thus LGRs account for predicted growth defects. (C) Mixture model, Z-transformation and empirically defined LGR cut-offs were compared for the Nop10 SPI dataset. Cut-off definition using a bimodal mixture model predicted more than twice the number of growth defects compared to Z-transformation or LGR-based thresholds.

## Discussion

ScreenGarden is a useful tool for easy, quick and robust analysis of plate-based high throughput assays and facilitates screen analysis that use two independent controls. Data can be plotted immediately without exporting output files into a second application for data visualisation, and ScreenGarden analysis does not require prior experience with handling of large-scale data. ScreenGarden is an open-source Shiny R application. All code is written using RStudio and available for download from the ScreenGarden homepage or GitHub. Thus, ScreenGarden can be run not only as a web application but also locally using the open source RStudio software, which runs on Windows, Mac and Linux platforms. This renders the possibility to adapt the code for screen-specific needs and easy customisation of the code. All ScreenGarden tools can be run independently, since the files are directly uploaded for each specific step. The normalisation and smoothing algorithms prevent biases due to plate differences or spatial anomalies, making ScreenGarden a robust tool for data analysis. ScreenGarden can perform analysis within seconds and provides data visualisation. Plots can be downloaded as PDF files for further preparation or directly incorporated into presentations or reports. We have shown that ScreenGarden analysis can identify reproducible growth defects at least as well as *ScreenMill*. Thus, ScreenGarden provides an easy to use software tool for plate-based microbial screen analysis.

## Resources

ScreenGarden can be run online as a Shiny R web application or locally using RStudio. ScreenGarden web-application: https://screengarden.shinyapps.io/screengardenapp/

ScreenGarden R scripts: https://github.com/CinziaK/ScreenGarden

If the user decides to run ScreenGarden locally, the following packages need to be installed prior ScreenGarden analysis:

tidyverse^21^, lubridate^22^, rlang, ggplot2^23^, Cairo, gghighlight, shiny, shinythemes, mclust^20^

