## Supplementary file 1 for "ScreenGarden: A shinyR application for fast and easy analysis of plate-based high-throughput screens"

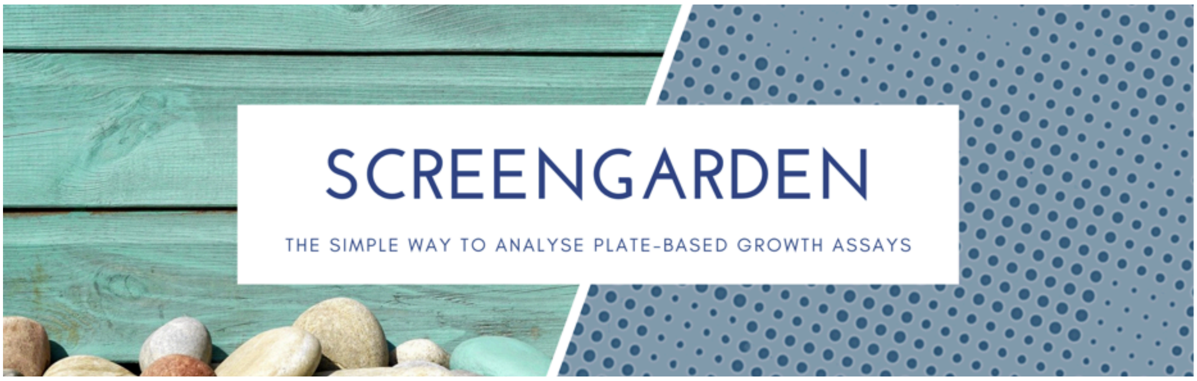


### A guide to data analysis using ScreenGarden

1. Input files

ScreenGarden requires a file which contains colony sizes on plate and a key-file, which contains information about the genotype for the specific plate position.

The key-file should be in .txt or .csv format and should be numeric for Plate, Row and Column values:

| Plate | Row | Column | ORF |
|---|---|---|---|
| 1 | 1 | 1 | YML032C |
| etc. | | | |

The colony-size file should be a .txt file and either derived from ScreenMill's CM Engine in the format:

| name,1,.tif | |
|---|---|
| 327 | 0.9381 |
| 271 | 0.9666 |
| 199 | 0.9356 |
| 195 | 0.9469 |
| 197 | 0.9262 |
| etc. | |

or converted to the format:

| Label | Plate | colonysize | Row | Column |
|---|---|---|---|---|
| name | 1 | 250 | 1 | 1 |
| etc. | | | | |

2. Calculate LGRs

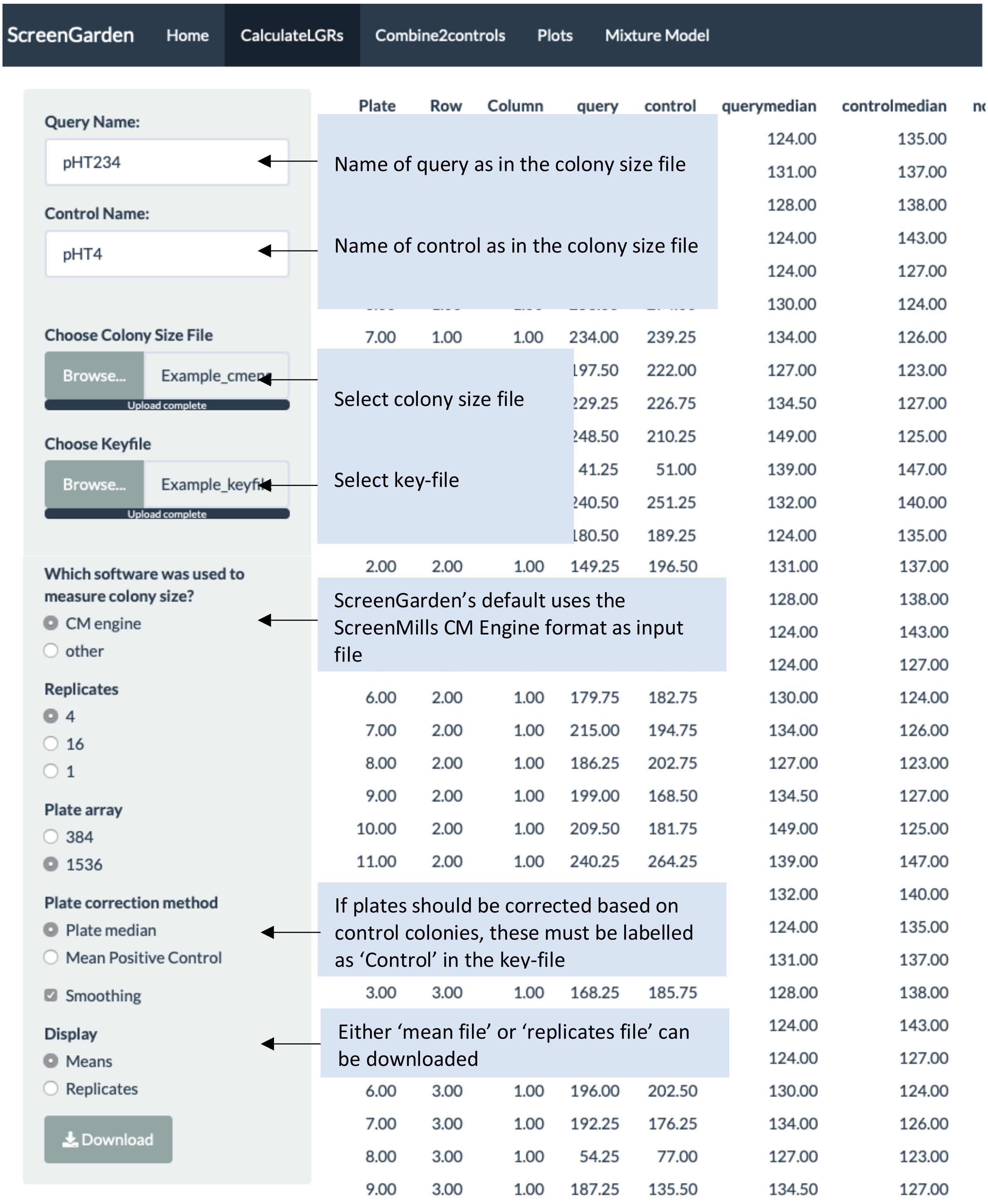


ScreenGarden is case-sensitive! Please make sure that query and control names are correct and the same as in the colony size file!

3. Combine2Controls

Two mean-file datasets can be combined to a merge file.

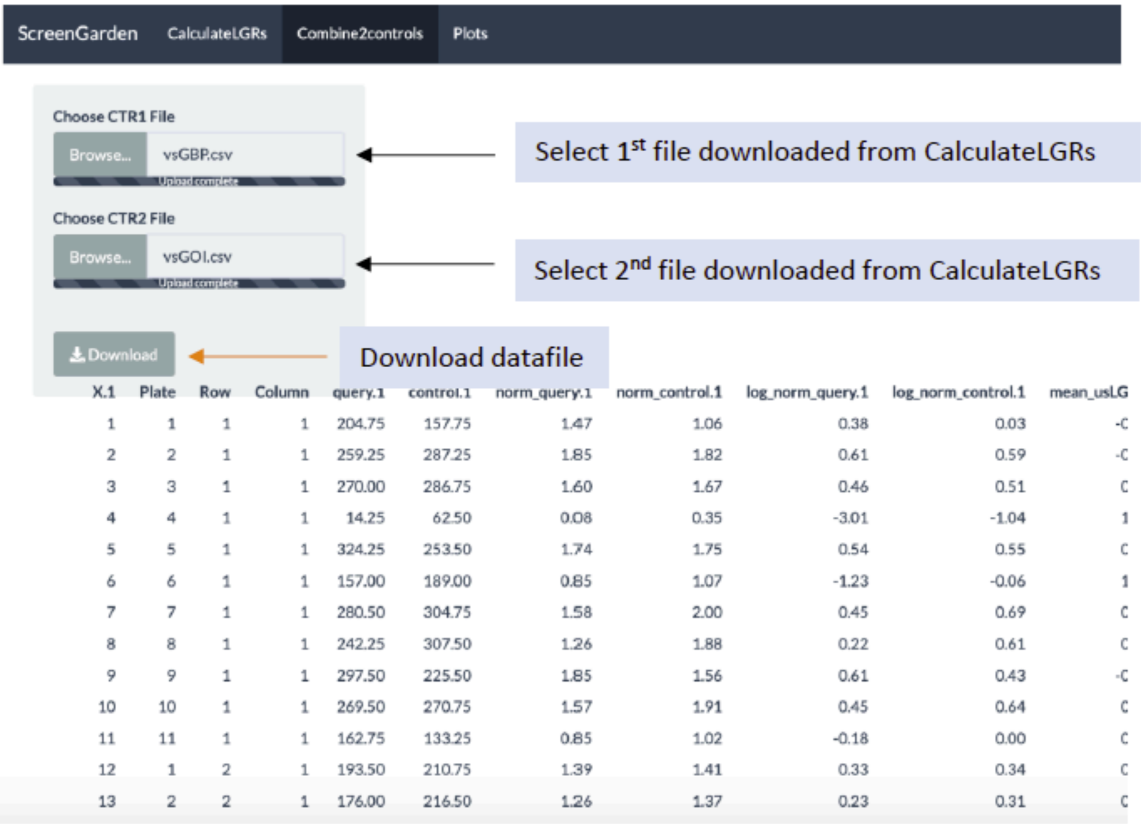


| X.1 | Plate | Row | Column | query.1 | control.1 | norm_query.1 | norm_control.1 | log_norm_query.1 | log_norm_control.1 | mean_usLG |
|---|---|---|---|---|---|---|---|---|---|---|
| 1 | 1 | 1 | 1 | 204.75 | 157.75 | 1.47 | 1.06 | 0.38 | 0.03 | -C |
| 2 | 2 | 1 | 1 | 259.25 | 287.25 | 1.85 | 1.82 | 0.61 | 0.59 | -C |
| 3 | 3 | 1 | 1 | 270.00 | 286.75 | 1.60 | 1.67 | 0.46 | 0.51 | C |
| 4 | 4 | 1 | 1 | 14.25 | 62.50 | 0.08 | 0.35 | -3.01 | -1.04 | 1 |
| 5 | 5 | 1 | 1 | 324.25 | 253.50 | 1.74 | 1.75 | 0.54 | 0.55 | C |
| 6 | 6 | 1 | 1 | 157.00 | 189.00 | 0.85 | 1.07 | -1.23 | -0.06 | 1 |
| 7 | 7 | 1 | 1 | 280.50 | 304.75 | 1.58 | 2.00 | 0.45 | 0.69 | C |
| 8 | 8 | 1 | 1 | 242.25 | 307.50 | 1.26 | 1.88 | 0.22 | 0.61 | C |
| 9 | 9 | 1 | 1 | 297.50 | 225.50 | 1.85 | 1.56 | 0.61 | 0.43 | -C |
| 10 | 10 | 1 | 1 | 269.50 | 270.75 | 1.57 | 1.91 | 0.45 | 0.64 | C |
| 11 | 11 | 1 | 1 | 162.75 | 133.25 | 0.85 | 1.02 | -0.18 | 0.00 | C |
| 12 | 1 | 2 | 1 | 193.50 | 210.75 | 1.39 | 1.41 | 0.33 | 0.34 | C |
| 13 | 2 | 2 | 1 | 176.00 | 216.50 | 1.26 | 1.37 | 0.23 | 0.31 | C |

4. Plotting data using ScreenGarden.

Mean or merge files can be uploaded and Columns can be plotted against each other. Plots can be downloaded directly from the website.

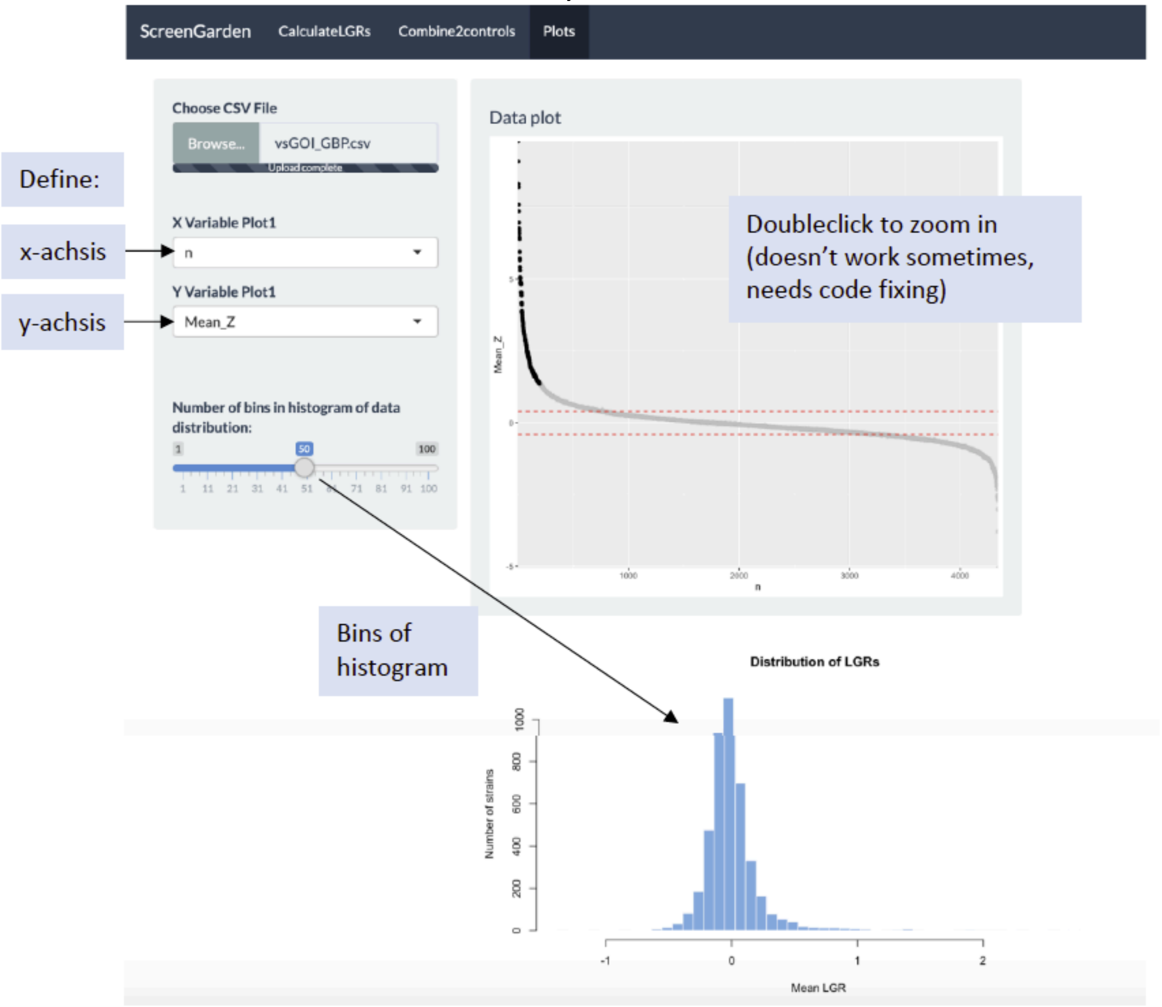
